# A synaptic threshold mechanism for computing escape decisions

**DOI:** 10.1101/272492

**Authors:** D.A Evans, A.V. Stempel, R. Vale, S. Ruehle, Y. Lefler, T. Branco

**Affiliations:** MRC Laboratory of Molecular Biology, Cambridge, CB2 0QH, UK; UCL Sainsbury Wellcome Centre for Neural Circuits and Behaviour, London, W1T 4JG, UK

## Abstract

Escaping from imminent danger is an instinctive behaviour fundamental for survival that requires classifying sensory stimuli as harmless or threatening. The absence of threat allows animals to forage for essential resources, but as the level of threat and potential for harm increases, they have to decide whether or not to seek safety^1^. Despite previous work on instinctive defensive behaviours in rodents^2–13^, little is known about how the brain computes the threat level for initiating escape. Here we show that the probability and vigour of escape in mice scale with the intensity of innate threats, and are well described by a theoretical model that computes the distance between threat level and an escape threshold. Calcium imaging and optogenetics in the midbrain of freely behaving mice show that the activity of excitatory VGluT2+ neurons in the deep layers of the medial superior colliculus (mSC) represents the threat stimulus intensity and is predictive of escape, whereas dorsal periaqueductal gray (dPAG) VGluT2^+^ neurons encode exclusively the escape choice and control escape vigour. We demonstrate a feed-forward monosynaptic excitatory connection from mSC to dPAG neurons that is weak and unreliable, yet necessary for escape behaviour, and which we suggest provides a synaptic threshold for dPAG activation and the initiation of escape. This threshold can be overcome by high mSC network activity because of short-term synaptic facilitation and recurrent excitation within the mSC, which amplifies and sustains synaptic drive to the dPAG. Thus, dPAG VGluT2^+^ neurons compute escape decisions and vigour using a synaptic mechanism to threshold threat information received from the mSC, and provide a biophysical model of how the brain performs a critical behavioural computation.

---

Detecting and escaping threats, such as predators or objects on collision course, is an instinctive animal behaviour that reduces the chances of being harmed or killed^12^. However, threats often co-exist with desirable resources, such as food or mates, and thus while escape ensures safety, it also results in halting other behaviours and a potential loss of resources^14^. To optimally balance escape with other survival behaviours, animals must use sensory information and past experience to estimate threat levels and decide whether or not to escape^1^, as well as how vigorous the defensive response should be^15–17^. While perceptual decision making has been extensively studied in primates and rodents using learned choice tasks^18–20^, and previous work has identified some of the key circuit nodes in the innate defensive system^4–8,13,21,22^, the neurophysiological basis of instinctive escape choices in mammals is largely unknown.

To investigate escape behaviour in mice we used overhead expanding spots, which are innately aversive as they mimic an approaching predator or object^3,23–25^, and varied the stimulus contrast to manipulate the probability of the stimulus being perceived as a threat. Animals were placed in an open corridor with a shelter at one end and a threat zone at the other, and stimuli were presented after mice entered the threat area spontaneously, resulting in escape towards the shelter (Fig. 1A-C). Stimulus-evoked escape responses were, however, variable and probabilistic. Decreasing the stimulus contrast resulted in a progressive increase in reaction time and reduction in escape probability, producing chronometric and psychometric curves qualitatively similar to those obtained in learned perceptual categorisation tasks^26^ (N=13 mice, 209 trials; Fig. 1D-E; Video 1; P=2.5×10^−7^ for escape probability, P=3.5×10^−8^ for reaction time, repeated measures ANOVA). In addition, the vigour of the response, quantified as the maximal escape speed, also increased gradually as a function of contrast (P=1.6×10^−6^, repeated measures ANOVA, Fig. 1F), showing that the probability, reaction time and vigour of instinctive escape are innately matched to the intensity of the threat stimulus (see also Fig. S1). We next modelled this relationship by modifying a single-layer network model previously applied to learned decision-making tasks^27^. This is a variant of drift-diffusion models^18,28^ that integrates a variable over time and implements the decision to escape as a threshold-crossing process. The key variable in the model is the threat level, which increases with sensory evidence of threat and decays over time (Fig. 1G, see Methods). The threat level is treated as a noisy decision variable, and escape is initiated when the variable crosses the escape threshold. In this model, the probability of escape and reaction time depend on the strength of sensory evidence and on the accumulated noise, and escape vigour is computed as a function of the peak threat level. This model produced a very good fit to the behavioural data obtained from visual stimulation (Fig. 1D-F). As a further test, we used innately aversive ultrasonic sweeps^29^, which generated escape with high probability, short reaction times and high vigour (Fig. 1 B-F), thus supporting a generic relationship between these variables.

Previous work has suggested a role for multiple brain regions in processing visually-evoked instinctive defensive behaviours^5,7,21,30^, so we next aimed to define circuit nodes that are critical for computing escape, using acute optogenetic inactivation with the chloride-conducting channelrhodopsin iChloC^31^ targeted to excitatory VGluT2^+^ neurons (Fig. S2A-B). Inactivation of the dPAG and mSC severely affected escape behaviour (Fig. 2A-B), but in different ways. Inactivation of the dPAG lead to a clear switch from escape to freezing in response to threats (P_escape_=3±3%, P_freeze_= 86±6%, mean freezing duration=4.3±1.0s; N=6 mice, P=8.12×10^−5^ for escape and P= 0.00029 for freezing, U-test for comparison between light-off and light-on; Video 2; Fig. 2A) with fast reaction times (269±35ms, Fig. 2A), indicating that the threat was still detected and a defensive action initiated, and that the dPAG is specifically required to initiate escape. On the other hand, both visual and sound stimuli after SC inactivation produced no defensive response in 62±10% of light-on trials, suggesting that the link between sensory stimulus and initiation of a response to threat was severely compromised (P_escape_=18±5%, P_freeze_= 19±7%, N=9 mice, P=5.15×10^−5^ for escape and P=0.02 for freezing U-test for comparison between light-off and light-on; Video 3). In the remaining trials, the reaction time was slow (1443±255ms, P=0.002, two-tailed t-test comparison with freezing reaction times after dPAG inactivation) and the vigour of escape low (77±7% of light-off, P=0.025 two-tailed paired t-test; Fig. S2C), compatible with a reduction in the perceived threat level. For both mSC and dPAG inactivation, there was no effect on speed during exploration, indicating that the effects on escape behaviour were not due to a non-specific disruption of motor function (Fig. S2D). Similar results were obtained with targeted muscimol inactivation of the dPAG and mSC, while in contrast, broad inactivation of regions such as visual cortex (V1) and amygdala lead to small decreases in response probability and vigour (Fig. S3), consistent with a modulatory role^30^. To further investigate the role of the dPAG and mSC in the computation of escape, we expressed the calcium indicator GCaMP6s in VGluT2^+^ neurons in the deep layers of the mSC (dmSC) or dPAG, and imaged calcium activity using head-mounted miniature microscopes^32^ in freely behaving animals during stimulus presentation. Neurons in both areas showed clear increases in calcium signals during stimulus-evoked escape trials (57 out of 138 dPAG cells from 3 mice, 55 trials; 177 out of 218 dmSC cells from 8 mice, 111 trials; Fig. 2C,F). Responding neurons had a mean reliability per trial of 28±3% for the dPAG and 35±3% for the dmSC, yielding a mean fraction of active cells per trial of 14±5% and 23±6%, respectively, which was stable over multiple trials (Fig. S4). However, the temporal profile of the activation of dPAG and dmSC was distinct. While dPAG cells were active in the peri-escape initiation period (mean onset= −0.24±0.21s, not different from 0s, the escape onset; P=0.24 two-tailed t-test; Fig. 2D,E), activity in the majority of dmSC cells preceded escape onset (mean onset= −1.51±0.17s, P=3.5×10^−12^ Wilcoxon test comparison with 0s, the escape onset; Fig. 2G,H). The distribution of dmSC activity onsets was bimodal, but analysis of single trials revealed that most cells did not respond exclusively in the pre- or peri-escape period (in 75% of cells the onset varied across trials between pre-and peri-escape), suggesting that the distribution does not reflect two distinct neuron populations with different roles, but rather a bias on the activity onset of individual cells. On each trial, the onset of the dmSC ensemble activity was −1.77±0.5s relative to the onset of escape (significantly different from escape onset, P= 0.00075, two-tailed t-test), whereas the onset of dPAG ensembles was −0.25±0.48s (P=0.59, two-tailed t-test comparison with escape onset), further confirming a temporal difference in the activation of these two networks.

**Figure 1.**
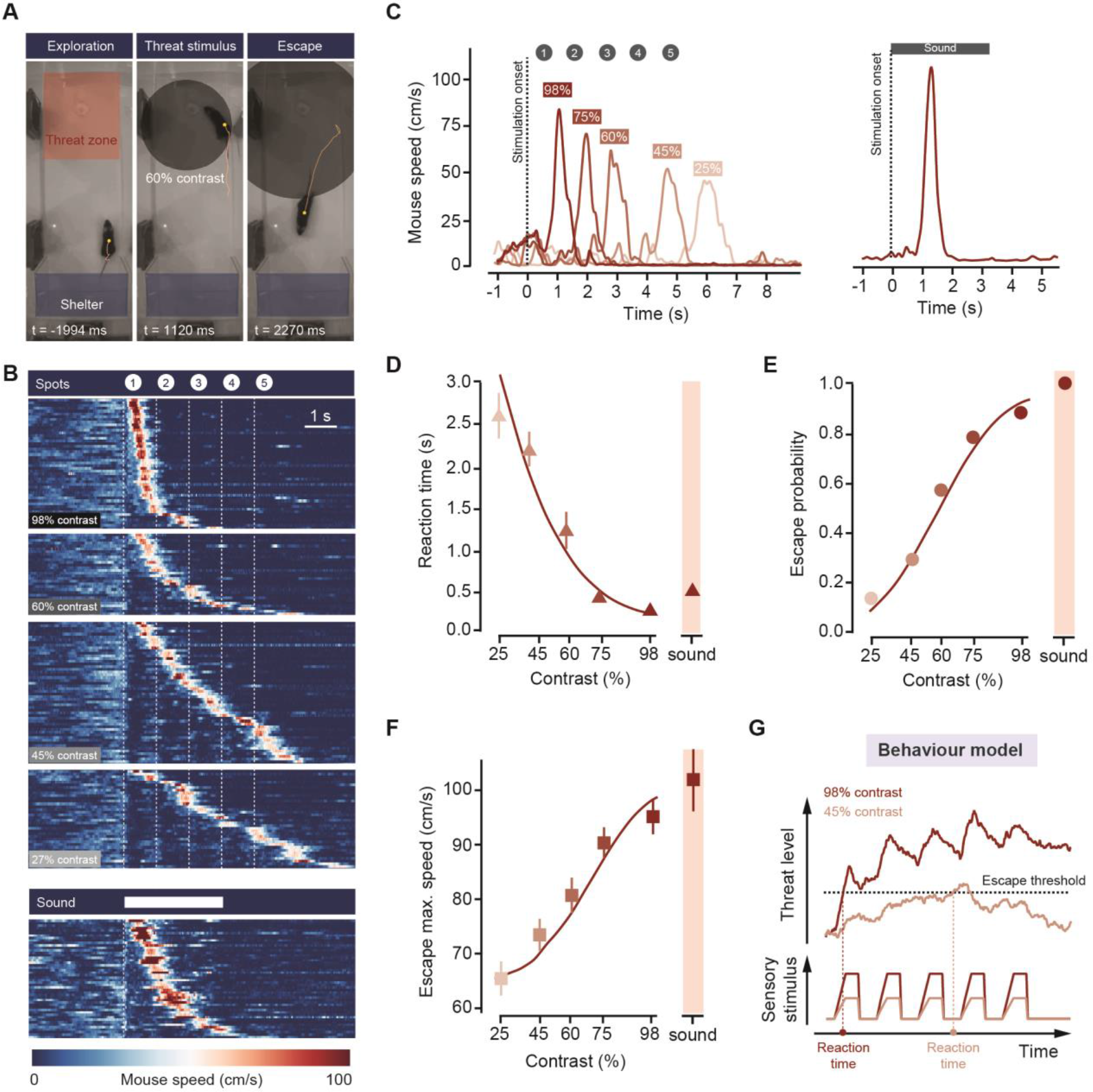
Escape behaviour during threats of varying intensity. **(A)** Video frames from one trial showing an escape response during stimulation with an expanding spot projected from above. Yellow lines indicate the mouse trajectory during the preceding 2 sec and frame times are relative to stimulus onset. **(B)** Raster plot of mouse speed for successful escape trials for visual (top, responses to 4 different contrasts shown) and sound (bottom) stimulation, from 13 animals, sorted by reaction time. **(C)** Example speed traces from one mouse in response to a train of spots at different contrasts (left) and to sound stimulation (right). Traces are from single trials. **(D)** Chronometric and **(E)** psychometric curve for stimulus intensity and escape behaviour, obtained by pooling data from all animals. Escape probability in **E** is for the first spot presentation (0-1140ms) **(F)** Summary plot from all animals showing that escape vigour increases as a function of stimulus intensity. **(G)** Schematic illustrating a model for computing escape behaviour threats of varying intensity. A sensory stimulus is integrated into threat level over time and escape is initiated if the level goes above the threshold for escape. The process is noisy, and as the threat level decreases the probability of crossing the threshold decreases. The sensory stimulus trace shows the diameter of the expanding spot scaled by the contrast. The threat level traces are example trials from simulations using the model parameters fit to the data. Datapoints in **D,E,F** are means for trials pooled across all animals (see also Fig. S1), error bars are SEM and continuous red line is the model fit to the data.

To determine whether dmSC and dPAG activity reflects the stimulus or the escape choice, we separated trials from the same stimulus intensity by trial outcome (Fig.2I). This analysis showed that dmSC neurons encode not only the presence of the threat stimulus, but also reflect the choice to escape (z-score=1.93±0.23 for escape, 1. 18±0.11 for no escape; P=0.023, two-tailed t-test between escape and no escape; P=5.8×10^−10^ 1-sample t-test between no escape and 0), whereas activity in dPAG neurons increases exclusively during escape trials (z-score=2.28±0.17 for escape, 0.49±0.19 for no escape; P=0.00028, two-tailed t-test between escape and no escape; P=0.11 1-sample t-test between no escape and for no escape; P=0.00028, two-tailed t-test between escape and no escape; P=0.11 1-sample t-test between no escape and 0). Receiver-operator characteristic analysis (ROC) of the ensemble activity reflected this difference and showed that the 5dPAG is an almost perfect classifier of the trial outcome (auc=0.92), while the dmSC is a noisier, but still very good classifier (auc=0.78). Further evaluation of the ROC evolution from stimulus onset showed that an ideal observer of dmSC activity could correctly predict the decision to escape above chance level 900ms before escape initiation (68% correct; Fig. 2J). In addition, the slope of the dmSC ensemble activity was strongly anti-correlated with the latency between stimulus presentation and escape (Pearson’s r=−0.94, P=0.0048), further suggesting that dmSC activity is an important determinant of escape initiation (Fig. 2K, Fig. S5A). To further test the nature of dmSC signals, we repeatedly exposed mice to high intensity threat stimuli, after which they developed strong place aversion (time spent in threat area 35.1±3.5% for naïve animals and 5.1±2.0% after conditioning, N=7 mice, P=2.2×10^−5^, two-tailed t-test, Fig. S6A-C) and initiated spontaneous flights upon approaching the threat presentation area (P_spontaneous escape_ 3.2±0.8% for naïve animals and 12.2±2% after conditioning, N=14 mice; P=0.004, two-tailed t-test; Fig. S6D; Video 4). Analysis of dmSC activity during conditioned flights showed a clear activity increase upon place entry and preceding escape, despite no stimulus presentation (mean z-score=1.94±0.17, n=57 trials from N=7 mice, P=0.00013, two-tailed t-test between pre and postconditioning; Fig. 2L). Importantly, pre-escape activity in these conditions was still predictive of escape (auc at escape onset = 0.74, significantly above chance 2.1s before escape), and not related to head-rotation movements (Fig. S5B), indicating that dmSC neurons encode a general variable that is correlated with the likelihood of escape. In agreement with the threat stimulus data, dPAG neurons were silent upon entering the threat area and only showed increased activity at escape initiation (Fig. 2L). Further analysis of the escape-evoked calcium signals showed a significant correlation between escape speed and peak calcium activity, which was ~3 times stronger in dPAG than in the dmSC (PAG: Pearson’s r=0.7, P=6.7×10^−7^; SC: r=0.25, P=0.04; Fig. 2M), and which was specific for running during escape to shelter and not present during exploration running bouts of similar speed (Fig. S5C).

**Figure 2.**
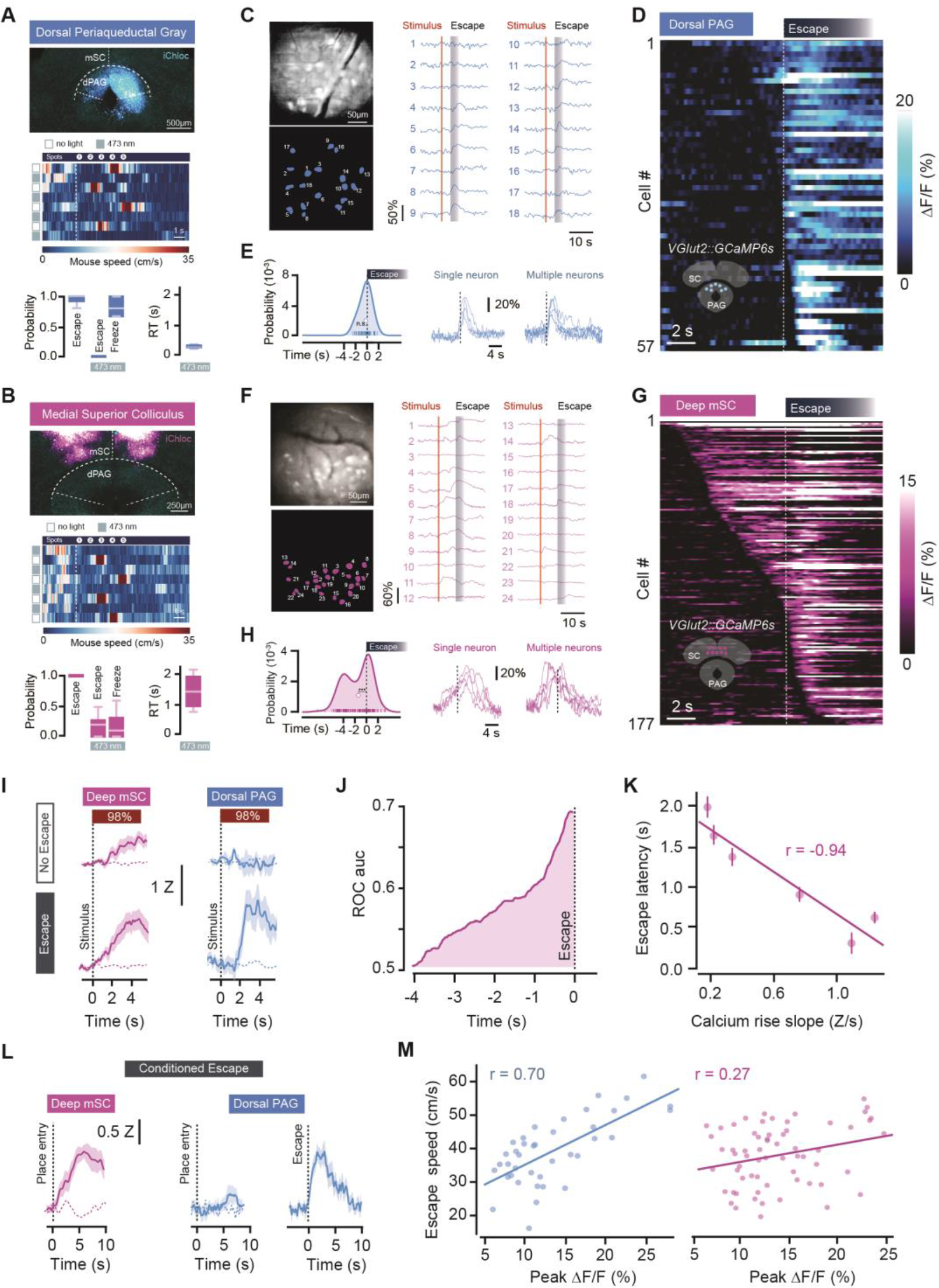
Encoding of threat and escape behaviour in the superior colliculus and periaqueductal gray. **(A)** Example image of iChloc expression in dPAG VGIuT2^+^ neurons (top), with example raster for mouse speed during interleaved trials of threat presentation with light off or on (middle), and summary data from visual and sound stimuli showing a switch from escape to freezing during dPAG inactivation (bottom, RT is reaction time). **(B)** Threat presentation after iChloc activation in the mSC strongly reduces the display of any defensive response. **(C)** Example GRIN lens field-of-view of GCamp6s in dPAG VGIuT2^+^ neurons (top left), with respective cell mask (bottom left) and single trial trace examples (right). **(D)** Raster showing the average calcium response for all recorded dPAG cells, aligned to escape onset and sorted by response onset. **(E)** Left, distribution of calcium response onsets for all dPAG cells, where curve is the kernel density estimation and markers are the onset of each cell (white marker shows the mean value). Right, single trial traces for a single neuron (left) and for multiple neurons in the same field of view (right). **(F,G,H)** Same as in (C,D,E) for dmSC cells. **(I)** Average population activity across all trials in response to 98% contrast spots (z-score), sorted by trial outcome for dPAG (blue) and dmSC (pink). Shaded areas are SEM and dashed lines are activity with no stimulus presentation. **(J)** Evolution of area under the curve from ROC analysis for dmSC signals up to escape onset. **(K)** Correlation between the rise slope of the population activity and escape latency. **(L)** Average population activity after conditioning with high intensity threats. Approaching the threat presentation area elicits an increase in dmSC population activity (left, dashed line is average activity before conditioning), whereas dPAG activity increases selectively upon escape, and not threat zone entry (middle and right; peak z-score for first 5s after threat area entry: 0.45±0.1 before conditioning (middle, dashed line) and 1.5±0.2 after conditioning, N=20 trials, P=0.0004, two-tailed t-test). **(M)** Correlation between the population activity of dPAG (left) and dmSC (right) and escape speed. Each data point is a single trial. Error bars and shaded areas in plots in all panels are SEM.

In the framework of our model, these activity profiles are consistent with dmSC neurons representing a preescape variable such as threat intensity, while dPAG neurons encode the result of the thresholding computation. This predicts that direct activation of dmSC neurons should produce psychometric and chronometric curves similar to sensory stimulation, as dmSC activity is still being passed through the threshold mechanism to initiate escape, while stimulation of dPAG neurons above the action potential firing threshold should reliably elicit escape behaviour with short reaction times. We tested this prediction by expressing Channelrhodopsin-2 (ChR2)^33^ in VGluT2^+^ neurons^34^ of dmSC or dPAG and delivering light stimulation *in vivo*^35^ (Fig. 3A). Light stimulation of both dmSC and dPAG recapitulated escape behaviour in response to threats, and elicited shelter-directed flights over a range of stimulation frequencies (Fig. S7A-B, Video 5). Next, we tested the behavioural effect of gradually increasing network activation by increasing light intensity^36^, which for dmSC neurons resulted in a progressive increase in the probability of escape while decreasing the variability of the responses (Fig.3B-C). In contrast, increasing activity in the dPAG network produced a steep, all-or-none curve, with stereotyped responses for each intensity (SC: N=278 trials from 4 mice; PAG: N=590 trials from 7 mice; Fig. 3B-C). Logistic regression confirmed that the slope of the dPAG psychometric curve was significantly steeper than for the dmSC (slope of logistic fit = 26.3, 95% CI [22.1, 30.4] for dPAG and 4.0, 95% CI [2.75, 5.25] for dmSC), in agreement with our model hypothesis. In addition, the latency to initiate escape decreased with stronger activation of dmSC neurons, while escape latencies for dPAG activation were short across the full stimulation intensity range (slope of linear fit for SC=-0.21, 95% CI [0.27,-0.15]; PAG=-0.07, 95% CI [-0.11,-0.03]; Fig. 3D), demonstrating that activity levels in the dmSC determine the onset of escape, as suggested by the calcium imaging data and predicted by the model. Increasing the stimulation strength of both networks was also correlated with an increase in escape speed, but the correlation was much stronger for dPAG stimulation (Pearson’s r=0.93, P=1.5×10^−5^) than for dmSC (Pearson’s r=0.58, P=0.04; Fig. 3E), which further supports a model where activity of dPAG neurons represents a post-threshold variable from which escape vigour is computed. Moreover, optogenetic activation of the dmSC while inactivating the dPAG with muscimol did not produce escape behaviour, whereas inactivation of an alternative dmSC projection target, the parabigeminal nucleus (PBGN)^5^, did not impair escape, suggesting that threat information from the dmSC has to flow through the dPAG in order to initiate escape (Fig. S8).

**Figure 3.**
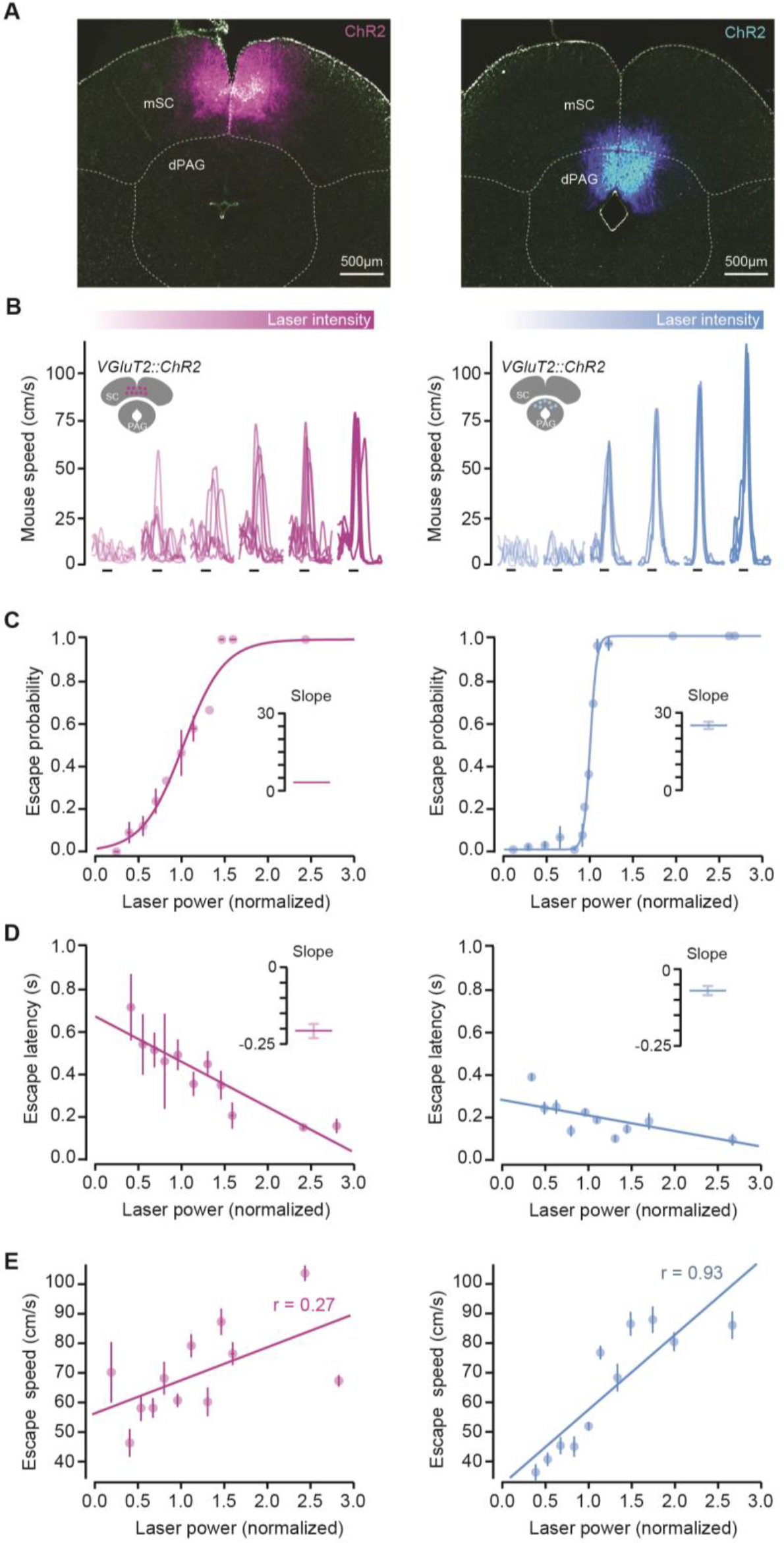
Optogenetic stimulation shows different roles for mSC and dPAG in escape behaviour. **(A)** Images of ChR2-EYFP expression in VGIuT2^+^ cells of mSC (left) and dPAG (right). **(B)** Example speed traces for stimulation with increasing blue light intensity at 10Hz (1s pulse train) for one mouse (dPAG right, dSC left). Each trace is from a single trial, black lines show stimulation timing. **(C)** Psychometric curve for light intensity versus escape probability showing a gradual increase for dSC stimulation (left) and an all-or-none profile for the dPAG (right). Lines are logistic fits to the data (pooled across all animals and binned light intensities), and insets show the slope of the fit (error bar is SD). **(D)** Increasing the stimulation light intensity causes a large reduction in escape latency for mSC (left), while the change in latency is much smaller for dPAG stimulation (right). Lines are linear fits to the data. Inset shows slope of linear fit (error bar is SD). **(E)** Correlation between light intensity and escape speed is stronger for dPAG stimulation (right) than for mSC (left). Error bars in all panels are SEM unless otherwise indicated.

To determine whether dPAG neurons receive information directly from the dmSC we first performed monosynaptic rabies tracing^37,38^ from starter dPAG VGluT2^+^ neurons. This resulted in extensive labelling of cells in the SC (Fig. 4A, Fig. S9A) in agreement with previous suggestions^39,40^, and revealed a 11:1 SC to dPAG convergence ratio, with the majority of the input cells being excitatory (87.9±1.0%, Fig. S9C-D) and distributed amongst the deep and intermediate SC layers. Moreover, most presynaptic SC cells were located in the medial part of the SC (82.9±2.6% of 1770 cells within medial bisection of ipsilateral SC, N=3; Fig. S9B) and dPAG neurons did not project back to the SC (N=3, Fig. S9E-F), indicating a feed-forward, columnar organisation of connectivity between the SC and dPAG. To investigate the properties of dmSC input to dPAG neurons, we used ChR2-assisted-circuit mapping by expressing ChR2 in VGluT2^+^ dmSC neurons and performed whole-cell patch-clamp recordings in acute midbrain slices. Light stimulation elicited excitatory monosynaptic input in 60% of VGluT2^+^ dPAG neurons (N=74 cells from 21 mice; Fig. 4B, left; see also Fig. S10), but the recorded connections were weak (peak EPSC: −37.9±11.9pA), exhibiting a high failure rate (20.3±8%) and a very low quantal content (2.3±0.6) even at full-field maximum light stimulation (Fig. 4B). Further analysis revealed that the statistics of neurotransmitter release were not significantly different from a Poisson model, indicating a very low synaptic release probability^41,42^ (slope of linear fit to direct quantal content versus log_e_(failure rate)^−1^ = 0.92, 95% CI [0.74,1.1]; Fig. 4C; Fig. S11A-C). An important consequence of such an unreliable connection is that the basal probability of eliciting action potentials in dPAG neurons from dmSC stimulation is extremely low (0.02±0.01 for single light pulses from resting membrane potential, N=29 cells; Fig. 4D), despite VGluT2^+^ dPAG neurons having a high input resistance (0.55±0.05GΩ, Fig. S11D-E), and thus provides a synaptic threshold for information represented in the dmSC to activate the dPAG escape network. However, repeated light stimulation at 10Hz and 20Hz elicited trains of action potentials (mean spikes/pulse: 0.17±0.1 for 10Hz and 0.16±0.08 for 20Hz), more than would be expected from temporal summation given that the inter-stimulus interval is ~2-3 times longer than the membrane time constant (τ=28.3±3ms, Fig. S11E, significantly different from the 20Hz inter-stimulus interval, P=5.8×10^−6^, 1-sample t-test against 50ms). This happens for two main reasons: first, the short-term plasticity dynamics of the connection are facilitating on average (20Hz PPR=1.22±0.09, 10Hz PPR=1.04±0.08), which is consistent with its low release probability^43,44^ and provides input amplification at the synaptic level (Fig. 4E). Second, dmSC stimulation triggered a large and long lasting increase in sEPSCs frequency (peak frequency change= 98±42Hz), that decayed to baseline with a 0.49s time constant (Fig. 4F). To determine the origin of this polysynaptic excitatory input, we recorded the connectivity between VGluT2^+^ dPAG-dPAG and dmSC-dmSC neurons. While we found weak synaptic input and sparse connectivity amongst dPAG cells (27%, −54±8.3pA, N=11 cells from 2 mice), 100% of dmSC cells received strong, polysynaptic input from other dmSC cells (−146.7±41.5pA, N=22 cells from 10 mice; Fig. 4G), in agreement with previous work^45^ and suggesting the presence of an excitatory recurrent network within the deep layers of the dmSC that provides signal amplification at the network level. Together, these synaptic and network mechanisms provide a means for sustained activation of the dmSC network to overcome the weak connection to VGluT2^+^ dPAG neurons and drive firing of the dPAG escape network. *In vivo* silicon probe recordings of single units in the dmSC of awake head-fixed animals^46^ (73 units from 3 animals, Fig. S12) next showed that in agreement with previous work^47,48^, during threat presentation, dmSC neurons fire action potentials in the short-term facilitation frequency range of the dmSC-dPAG synaptic connection, and in a contrast-dependent manner (peak firing rate during stimulus period: 20.4±4.1Hz for 98%, 10.7±1.8Hz for 50%, N=32 visual-responsive units; 23.9±2.5Hz for sound, N=45 sound-responsive units; P=0.01 for 50% vs 98% and P=2.8×10^−5^ for 50% visual vs sound, U-test; Fig.4H). Moreover, a fraction of the stimulus-responsive units showed increased firing above baseline that persisted beyond the stimulus duration (37% of visual-responding and 15% of sound-responding units; time constant to decrease to baseline: 5.8s for sound stimuli, and 0.23s for visual stimuli; Fig. 4I), in agreement with our model that threat stimuli can engage a recurrent dmSC network that assists with integration to threshold.

**Figure 4.**
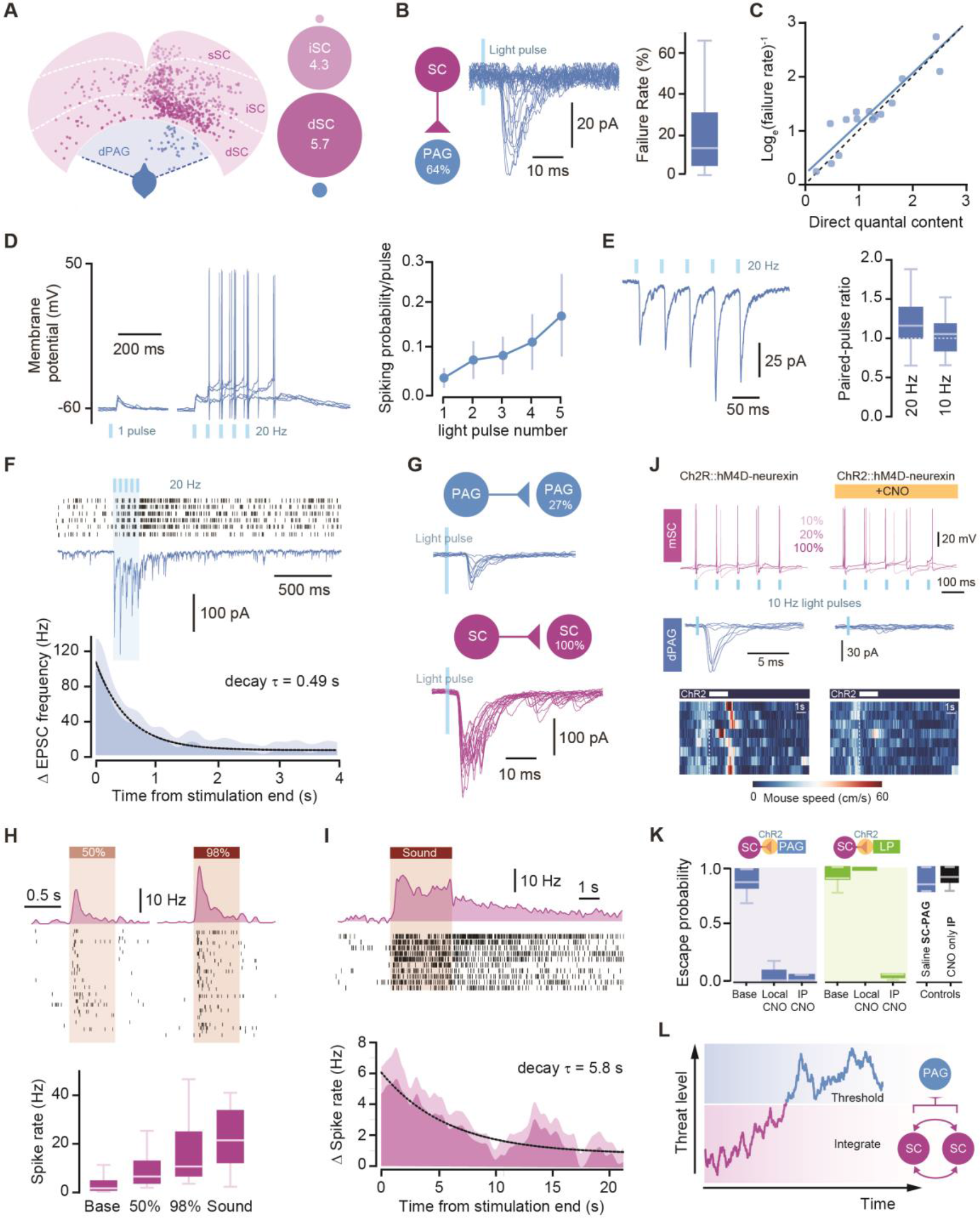
Neural circuit and biophysical mechanisms for computing escape behaviour. **(A)** Left, schematic illustrating the position of starter dPAG VGIuT2^+^ (blue) and presynaptic SC cells (pink) labelled via monosynaptic rabies tracing, for deep, intermediate and superficial SC layers (dSC, iSC, sSC respectively, see also Fig. S9). Right, ratio of presynaptic SC cells to PAG cells, depicted by circle areas (blue circle is one PAG cell, numbers show SC:PAG cell ratios). **(B)** Schematic of connectivity rate (left) and single traces for ChR2-evoked mSC-dPAG EPSCs (middle). Right, summary failure rate of mSC-dPAG excitatory connections. **(C)** Plot of direct quantal content versus estimation from failure rate assuming Poisson statistics. Blue line is linear fit to the data, dashes are unity line. **(D)** Membrane potential in response to single and 20Hz light stimulation (left, traces are individual trials), and respective summary quantification of spiking probability (right). **(E)** Average trace for one cell showing short-term facilitation at 20Hz stimulation (left) and quantification for all cells (right). **(F)** Example voltage-clamp trace for a dPAG cell single trial of 20Hz mSC stimulation (middle) and raster of sEPSCs for 5 trials in the same cell (top). Bottom, summary histogram of the increase in sEPSCs in dPAG cells with 20Hz mSC stimulation. Dashed line is single exponential fit. **(G)** Schematics of connectivity rate between dPAG (top) and dSC neurons (bottom), and respective example traces for single cells. **(H)** Top, example firing rate histograms and respective raster plots of consecutive trials for one single unit in the dmSC responding to visual stimuli of low (left) and high contrast (right). Bottom, summary quantification of average spike rate during baseline, and peak spike rate during threat stimuli. **(I)** Top, example unit showing persistent activity after sound stimulation, and average histogram for all cells with persistent activity after sound presentation (bottom, dashed line is single exponential fit). **(J)** Example *in vitro* recordings showing that hM4d-neurexin activation with CNO does not affect ChR2-evoked action potentials in the mSC (top, pink numbers are light intensity), but blocks mSC-dPAG EPSCs (middle). Bottom, example behaviour raster plots of multiple trials of mSC optogenetic activation showing that blocking the mSC-dPAG projection abolishes escape. **(K)** Summary for ChR2::hM4D-neurexin data showing that escape behaviour is selectively blocked by perturbing mSC-dPAG synapses with CNO local microinfusions, but not mSC-LP connections. Saline mSC-dPAG infusion and CNO i.p. in the absence of hM4D does not reduce escape (P>0.15, U-test, N=5 mice) **(L)** Schematic of a model for computing escape decisions. Threat evidence is integrated by a recurrent excitatory network of mSC neurons, and the mSC-dPAG connection sets the threshold for initiating escape. When activity in the mSC network is high enough, it drives firing of dPAG neurons, which represent the result of the thresholding computation and determines escape initiation and vigour. Shaded areas in histograms show SEM.

Given that the dmSC projects to several downstream targets^49^ and that the dPAG receives multiple afferents^50^, in the final set of experiments we used a chemogenetic approach to test whether the dmSC-dPAG synaptic connection is a critical pathway for computing escape from threat. We co-expressed the synaptically-targeted inhibitory designer receptor hM4D-neurexin^51^ and ChR2 in VGluT2^+^ dmSC neurons, which caused a 71±7% reduction in synaptic transmission to the dPAG in the presence of clozapine-N-oxide (CNO), while leaving firing in dmSC neurons intact (Fig. 4K, Fig. S13 A-C). *In vivo* microinfusion of CNO over dmSC-dPAG synapses lead to an almost complete block of escape behaviour to both visual (Fig. S13B, Video 6) and optogenetic activation of the dmSC, of similar magnitude to systemic injection of CNO (P_escape_=8±5% after local CNO for N=5 mice, P=0.0008 U-test test against baseline escape probability; P_escape_=5±5% after i.p. CNO for N=4 mice, P=0.01 U-test test against baseline escape probability; P=0.5 for local vs i.p. CNO U-test; Fig. 4K-L). Notably, doubling the intensity or frequency of optogenetic stimulation, which increases firing of dmSC neurons by 40% *in vitro* (Fig.S13A), was not sufficient to rescue escape behaviour (Fig. S13D), and applying the same strategy to inhibit the dmSC projection to the lateral posterior nucleus of the thalamus (LP), which has been shown to be involved in freezing behaviour, did not affect escape (P_escape_=100±0% after local CNO for N=4 mice, P=0.1 U-test test against baseline escape probability; P_escape_=4±2% after i.p. CNO for N=3 mice, P=0.04 U-test test against baseline escape probability; P=0.01 for local vs i.p. CNO U-test; Fig. 4L).

Our results are compatible with a two-stage model where evidence about threats is integrated in the dmSC network and passed through a threshold at the dPAG level to initiate defensive escape. We suggest that an important component of this threshold is the low release probability connection between VGluT2^+^ dmSC and dPAG neurons, which threats of high intensity have a higher probability of overcoming. Our data support a critical role of dPAG excitatory neurons in the computation of escape initiation and vigour, which is in agreement with work in several animal species, including humans, indicating that the PAG is a critical node in active defense^6,22,40,52,53,50,54–56^. While the mSC projects to several brain areas, and it is likely that different node projections contribute to the organisation of the complete defensive response, our inactivation experiments show that the dmSC-dPAG synaptic connection is necessary for escape initiation, whereas SC projection targets such as the PBGN or the LP^5,7^ are not. Given the proposed role of the mSC-LP projection in freezing behaviour^7^, our results suggest that there might be dedicated mSC projections for the control of freezing and escape. Optogenetic activation of the PBGN has been previously shown to elicit escape-like behaviour^5^, which in light of our results could be explained by antidromic activation of SC neurons that project to both the PBGN and dPAG, or by PBGN neurons projecting back to the SC. A key result of our study is that dmSC activity upon threat presentation is not a simple representation of the sensory stimulus, but encodes a higher order signal that is predictive of escape, such as the perceived threat level as in our behaviour model. The SC has a well-described role in multisensory integration^57,58^, and in our paradigm, dmSC neurons likely receive sensory input from the superficial SC for visual stimuli^30,58–60^ and from the inferior colliculus and auditory cortex for sound stimuli^8^. This higher-level computation in the SC is in agreement with previous work showing that the SC is involved in decision-making in primates and rodents^61–63^, and importantly, with results from human studies suggesting that the SC is part of an innate alarm system that detects and processes subliminal threat evidence^64,65^. Our demonstration of increased dmSC activity after a conditioning paradigm further supports a general role for the SC in processing generic threat cues, both innate and learned. Successfully escaping from threats to reach safety requires integration of multiple information streams, such as knowledge about the spatial environment^9^, and our results provide a mechanistic entry point for understanding how the brain computes a fundamental survival behaviour, and goal-directed behaviours in general.

## Experimental Procedures

**Animals**: Male and female adult C57BL/6J wild-type, VGluT2-ires-Cre (Jackson Laboratory, stock #016963) and VGluT2::EYFP (R26 EYFP, Jackson Laboratory #006148) mice were housed with ad libitum access to chow and water on a 12h light cycle and tested during the light phase. All experiments were performed under the UK Animals (Scientific Procedures) Act of 1986 (PPL 70/7652) following local ethical approval.

**Surgical procedures**: Animals were anaesthetised with an intraperitoneal injection (i.p.) of ketamine (95mg/kg) and xylazine (15.2mg/kg), and carpofen (5 mg/kg) was administered subcutaneously. Isoflurane (0.5-2.5% in oxygen, 1L/min) was used to maintain anaesthesia. Craniotomies were made using a 0.5mm burr and viral vectors were delivered using pulled glass pipettes (10μ1 Wiretrol II with a Sutter P-1000) in an injection system coupled to a hydraulic micromanipulator (MO-10, Narishige) on a stereotaxic frame (Model 1900 and 963, Kopf Instruments), at ~10nl/min. Implants were affixed using light-cured dental cement (RelyX Unicem 2, 3M) and the wound sutured (6-0, Vicryl Rapide) or glued (Vetbond). Coordinates are measured from lambda.

**Viruses**: The following viruses were used in this study and are referred to by contractions in the text. For optogenetic activation, AAV2-EF1a-DIO-hChR2(H134R)-EYFP-WPRE (3.9×10^12^ GC/ml), AAV2-EF1a-DIO-hChR2(H134R)-mCherry-WPRE (6.6×10^12^ GC/ml; Deisseroth) were acquired from the UNC Vector Core (USA). Optogenetic inhibition experiments were performed with AAV9-Ef1a-DIO-iChlo-2A-tDimer (3.75×10^12^ GC/ml; a gift from S. Wiegert and T. Oertner) or AAV1-EF1a-DIO-iChloc-2A-dsRed (5×10^13^ GC/ml; Addgene #70762, a gift from T. Margrie). For control and calcium imaging experiments respectively, AAV2-EF1a-DIO-EYFP-WPRE (4.0×10^12^ GC/ml) and AAV9-CAG-DIO-GCaMP6s-WPRE (6.25×10^12^ GC/ml) were acquired from Penn Vector Core (USA). For retrograde rabies tracing, EnvA pseudotyped SADB19 rabies virus (EnvA-dG-RV-mCherry) was used in combination with AAV8 coding for TVA and rabies virus glycoprotein (RG) that were prepared from pAAV-EF1a-FLEX-GT (Addgene plasmid #26198, Callaway) and pAAV-Syn-Flex-RG-Cerulean (Addgene plasmid #98221, Margrie). All viruses used for rabies tracing were a gift from T. Margie^66^. Additionally, a recombinant AAV with retrograde functionality (rAAV2-retro-mCherry, 6.97×10^12^ GC/ml, Addgene #81070^67^) was used. For chemogenetic inhibition experiments, AAV5-CAG-DIO-mCherry-2A-hM4D-HA-2A-nrxn1A (3.9×10^12^ GC/ml, a gift from S. Sternson) or AAV2-CAG-DIO-mCherry-2a-hM4D-nrxn1a (6.19×10^n^ GC/ml, Addgene #60544) were used.

### Behavioural Procedures

***Experimental set-up***: All behavioural experiments were performed in a rectangular perspex arena (W:20cm × L: 60cm × H: 40cm) with a red-tinted shelter (19cm × 10cm × 13.5cm) at one end, housed within a sound-deadening, light-proofed cabinet with six infra-red LED illuminators (TV6700, Abus). A screen (90cm × 70cm; ‘100 micron drafting film’, Elmstock,) was suspended 64cm above the arena floor, and a DLP projector (IN3126, InFocus) back-projected a grey uniform background via a mirror, providing 7-8lx at the arena floor. Experiments were recorded at 50 frames per second with a near-IR GigE camera (acA1300-60gmNIR, Basler) positioned above the arena centre. Video recording, sensory and optogenetic stimulation was controlled with custom software written in LabVIEW (2015 64-bit, National Instruments). The position of the animal was tracked on-line, and used to deliver stimuli when the animal entered a predefined ‘threat area’ (21cm × 20cm area at opposite end to shelter). An empty plastic Petri dish (replaced fresh for each animal; 35 mm) was affixed to the arena floor in the centre of the threat area to enrich the environment. All signals and stimuli, including each camera frame, were triggered and synchronised using hardware-time signals controlled with a PCIe-6351 board (National Instruments).

***Protocols***: Mice were placed in the arena and given 8min to explore the new environment, after which sensory stimuli were delivered when the animal entered the threat area longer than 100ms. A typical experiment lasted 30-60min. In the standard visual stimulation protocol, we used a pseudo-random contrast sequence to minimise the development of aversion or habituation during the behavioural session (see Fig.S6E-F for quantification). The sequence consisted of a first stimulus at 98% contrast, followed by a random selection without replacement from the remaining contrasts, and this process was repeated until the end of the behavioural session. Each stimulus was delivered with at least 30s inter-stimulus interval. For the conditioning protocol shown in Figures 2J and S4, repeated presentations (3-6 trials) at 98% contrast were delivered with no minimum inter-stimulus interval after a 10min acclimatisation period.

***Sensory stimuli***: The standard visual stimulus was a sequence of five dark expanding circles, and unless otherwise stated, each subtended a visual angle of 2.6° at onset and expanded linearly at 118°/s to 47° over 380ms, after which it maintained the same size for 250ms and began an inter-stimulus interval of 500ms. The contrast of the spot was varied in a number of experiments, and for clarity is reported as a positive percentage (low to high; e.g. 20% to 98%), converted from the negative Weber fraction (low to high; −0.2 to −0.98). The contrast was varied by altering the intensity of the spot against a grey screen maintained at constant luminance (standard luminance, 7.95cd/m^2^). The spot was located on the screen directly above the centre of the threat area, ~15° from the animals’ zenith. The auditory stimulus consisted of a frequency-modulated upsweep from 17 to 20kHz over 3s (Mongeau et al. (2003)). Waveform files were created in MATLAB (Mathworks), and the sound was generated in LabVIEW, amplified and delivered via an ultrasound speaker (L60, Pettersson) positioned 50-55cm above the arena, centred over the threat area.

***Analysis***: Behavioural video and tracking data was sorted into peri-stimulus trials and manually annotated. Detection of the threat stimulus was assumed if the animal showed a stimulus-detection response, in which the ears of the animal move posteriorly and ventrally, which precedes interruption or commencement of body movement. To differentiate failures of escaping from failures of attending to the stimulus, trials with no stimulus-detection response were excluded from the analysis. This resulted in the exclusion of three no-escape trials from the 25% contrast dataset, which increased the escape probability from 0.12 to 0.13. The onset of escape was measured as the first video frame marking the onset of a continuous movement consisting of a head turn followed by a full-body turn towards the shelter. Escape was annotated automatically and defined as the animal moving to enter the shelter in a single movement without stopping, within 0.9s after stimulus termination (or 6s after approaching a 15cm boundary from threat area for spontaneous escapes after conditioning). Behaviour metrics were calculated by pooling all trials and animals (Fig. 1D-F) and also by analysing each mouse individually and then computing an average value across all mice (Fig. S1A-C). Statistical analysis was performed using animal numbers for sample size. The escape probability for a given stimulus is the fraction of trials which led to an escape to the shelter. The maximum speed of the escape is calculated as the peak value of the speed trace between the onset of the escape and entry to the nest. Quantification of exploratory behaviour was done for behavioural sessions lasting at least 40 min, by calculating the cumulative displacement of the animal in 1 min bins followed by smoothing with a 5 point flat window. We did not observe any differences in the behavioural response to threat stimulation between male and female mice, and therefore data from both sexes has been pooled (for 98% contrast stimulation, escape probability: 0.86 for males, 0.88 for females, P=1.0, Fisher exact test; reaction time: 369.2±51.8ms for males, 365.6±39.6ms for females, P = 0.96, two-tailed t-test; vigour: 91.8±4.5cm/s for male, 89.1±11.1 for female, P=0.81, two-tailed t-test).

### Behavioural Model

The threat level (*T*) evolves over time according to

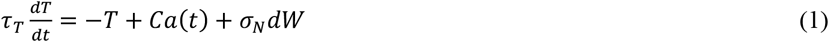

where *a*(t) is the diameter of the expanding visual spot scaled by the spot contrast *C*. The variable τ_T_ sets the time constant for changing the threat level and *dW* is a white-noise Wiener process parametrised by σ_N_. At each time point, *T* is compared against a threshold *B*, and escape initiated if *T* > *B*. The reaction time is the time at threshold crossing measured relative to stimulus onset. In this model we allow the threat level to continue evolving after the threshold has been crossed, similar to previous work on changes of mind during decision making^68^, and escape vigour *V* is computed from the peak of the threat level as a logistic function:

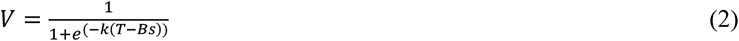

The model was first fitted with three free parameters (*B*, τ_T_, σ_N_) to the reaction time and escape probability data simultaneously by simulating 10,000 trials for each parameter set and using the brute force method in LMFIT Python 2.7 package. Escape vigour was then fitted to the average peak threat levels across all trials with free parameters *k* and *s* using least-squares minimisation in LMFIT. The fit parameters for the curves shown in Figure 1 are: *B*=0.165, τ_T_=1200ms, σ_N_=0.6, *k*=90, *s*=1.5.

### Pharmacological Inactivation

Animals were bilaterally implanted with guide cannulae (Plastics One, Bilaney Consultants) over the target region (see Supplementary Table 1) and given at least 48h for recovery. On test day, mice were placed in the standard arena for 10min and escape responses were assessed with a single visual stimulus (one 98% contrast expanding spot) or auditory stimulus. Additionally, in PBG- and PAG-cannulated mSC-VGluT2::ChR2 animals, optogenetic responses were also evoked. The animals were then lightly anaesthetised in an induction chamber and placed on a heating pad where anaesthesia was maintained with a nose-cone (2% isoflurane, 1L/min). Internal cannulae were inserted and sealed with Kwik-Sil. Muscimol-BODIPY-TMR-X (0.5mg/ml) or Alexa-555 (100μM; Life Technologies), dissolved in 1:1 PBS: 0.9% saline with 1% DMSO, was then infused at a rate of 70-100nl/min using a microinjection unit (10μl Model 1701 syringe; Hamilton, in unit Model 5000; Kopf Instruments) followed by a 5min wait period per hemisphere. Animals spent no longer than 30mins under anaesthesia and were given 30min to recover in the homecage, after which they were placed back in the cleaned arena and subjected to visual, auditory or optogenetic stimulation. Immediately upon termination of the behavioural assay, ~1hr after infusion, animals were anesthetised with isoflurane (5%, 2L/min) and decapitated. Acute slices (150μm) were cut using a microtome (Campden 7000smz-2 or Leica VT1200S) in ice-cold PBS (0.1M), directly transferred to 4% PFA solution, and kept for 20min at 4°C. The slices were then rinsed in PBS, counter-stained with DAPI (3μM in PBS), and mounted on slides in SlowFade Gold (Life Technologies) before wide-field imaging (Nikon TE2000) on the same day to confirm the site of infusion. Behavioural data was annotated as described. For the calculation of the maximum exploration speed, the peak speed of the 7min acclimatisation period before stimulation was used. Statistical analysis was performed using animal numbers for sample size.

### Calcium imaging in freely-moving animals

***Data acquisition***: A miniaturised head-mounted fluorescence microscope (Model L, Doric Lenses Inc.) was used to image GCaMP6s in neurons of male VGluT2-Cre mice. AAV9-CAG-Flex-GCaMP6s (300-550nl; Penn Vector Core) was injected into the mSC (AP: −0.2 to −0.5, ML: +0.25, DV: −1.6) or dPAG (AP: −0.4 to −0.6, ML:+0.25, DV: −2.2). At the level of the inferior colliculus, the dura was incised using a 30G needle, and gently pulled forward to partially reveal the SC. A GRIN lens-equipped cannula (SICL_V_500_80; Doric Lenses Inc.) was used to push forward the transverse sinus and inserted to the same depth as the injection coordinates, after which the craniotomy was covered with Kwik-Cast and the cannula affixed with dental cement. At least 21 days after surgery, the microscope was attached to the mouse without anaesthesia, and the animal was placed back in the homecage for 5-10min, for acclimatisation to the microscope. During this period, the optimal imaging parameters for the preparation were determined: the acquisition rate was 14.2Hz in most experiments (median; range: 10-20Hz) with an excitation power of 450 μW (median; range: 0.2-1.1mW). After a baseline period of 7min, animals were exposed to visual and/or auditory stimulation. For the visual stimulation, contrast was 98%, the inter-stimulus interval was 750ms, and post expansion period was 20ms, with the total epoch length and expansion rate unchanged. A typical session lasted 1.5hr (1-3 sessions per animal), with imaging data acquired during stimulation and control trials in ~5min epochs, with at least 2 days between sessions. If prolonged bouts of animal inactivity occurred, imaging was halted to minimize photobleaching. Fluorescence and behavioural frame trigger signals were acquired at 10kHz for offline synchronisation.

***Data analysis***: Behavioural video and tracking data were sorted into peri-stimulus trials and manually annotated to mark behavioural events as described above. Fluorescence stacks were registered^69^ and background-subtracted (Fiji). Cell body-like structures were identified manually as regions-of-interest (ROIs; elliptic or polygonal areas) in Fiji using the maximum intensity projection of registered movies, aided by inspection of deconvolved images. For each animal, ROI masks were rigidly translated to account for FOV movement between imaging sessions, and new cells added to the FOV if they became visible. In some cases, the FOV moved such that ROIs could not be mapped to the previous sessions, and it was therefore counted as a new FOV. Mean intensity traces were extracted for each ROI, interpolated with the behavioural video frames and tracking data, and ΔF/f calculated on a trial-by-trial basis with a baseline of 5s before stimulus onset. Traces were then smoothed with a 20 point Hanning window and Z-scored. ROIs were only included in the analysis if they had transients with a Z-score above 2 at any time during the recording session, to ensure that they were live, active neurons. Average responses for each cell were obtained by averaging across all trials independent of the trial outcome and statistical analysis was performed on all cells pooled together. Ensemble average responses were obtained by averaging the responses of all cells in a field-of-view and summary statistics calculated over all trials for each field-of-view. For the ROC analysis, the annotated behavioural outcomes were used to sort data into ‘Escape’ and ‘No Escape’ classes, and the ROC curves and AUC statistics were calculated using the open-source package Scikit-learn. The SD for the AUC was estimated using bootstrapping. ‘Peri’ and ‘Pre-escape’ time periods were defined as escape onset ±1s and <1s, respectively. For the plot in Figure 2K, escape latencies were first binned and average calcium signal waveforms calculated for each bin, and the signal rise slope was obtained by fitting a linear function (y=mx+b). The onset of calcium signals was measured by finding the time of the peak and iteratively moving backwards along the signal to determine the time point at which the signal reaches the baseline.

### Optogenetic experiments

For optogenetic activation, VGluT2-Cre and VGluT2::EYFP mice were injected with AAV-DIO-ChR2-EYFP or - mCherry, (see Viruses) into the dmSC (80-120nl per side, ML: +/− 0.2 to 0.35, AP: −0.25 to −0.45, DV: −1.4 to −1.55) or dPAG (40-80nl per side ML: +/− 0.0 to −0.4, AP: −0.4 to −0.6, DV: −1.95 to −2.2). Control animals were injected with 120nl AAV2-DIO-EYFP into the dPAG. One optic fibre (200 μm diameter, MFC-SMR; Doric Lenses Inc.) was implanted per animal, medially, 250-300 μm dorsal to the injection site. For optical stimulation, light was delivered by a 473nm solid-state laser (CNI) in conjunction with a continuous ND filter wheel for varying light intensity (NDC-50C-4M, Thorlabs) and a shutter (LS6, Uniblitz) driven by trains of pulses generated in LabVIEW. In some experiments, this system was substituted by a laser diode module (Stradus, Vortran) with direct analogue modulation of laser intensity. Magnetic patchcords (Doric Lenses Inc.) were combined with a rotary joint (FRJ 1×1, Doric Lenses Inc.) to allow the cannula to be connected without restraint and allow unhindered movement. In all experiments, animals were placed in the standard arena and given 8min to acclimatise. For the intensity modulation assay, the laser intensity was set initially to give a low irradiance (0.1-0.2mW/mm^2^) that did not evoke an observable behavioural response. Mice were photostimulated (473nm, train of 10 light pulses of 10ms at 10Hz) upon entering the threat area with an inter-stimulus interval of at least 30s. After at least three trials of this intensity, the irradiance was increased by 0.1-0.3mW/mm^2^ until a behavioural response was observed, after which 8-15 trials were obtained at a given intensity, before further increasing the light intensity. This process was iterated until an intensity was reached which always evoked a flight response (P_escape_=1). For one animal, the standard stimulus was not sufficient to reach P_escape_=1 and the curve was acquired with a higher frequency stimulus (10 light pulses of 10ms at 20Hz). If the animal stopped exploring the arena, precluding P_escape_=1 from being obtained, the experiment was terminated after 4hrs and not analysed. To normalise stimulation intensity and compare across animals, trials were first classified as escape if the animal reached the shelter within 5s of stimulation onset, to calculate the fraction of escape trials at a given intensity. The escape probability curve of each animal was then fitted with a logistic function (1/(1+*e*^−*k*(*x-x*0)^), and light intensities were normalised to *x*0. In the frequency modulation assay, high laser power was used (range, 12-13.5mW/mm^2^) and the stimulus consisted of 10 light pulses of 10ms at either 2, 5, 10, 20 and 40Hz, delivered in a pseudo-random order.

For histological confirmation of the injection site, animals were anaesthetised with Euthatal (0.15-0.2ml and transcardially perfused with 10ml of ice-cold phosphate-buffered saline (PBS) with heparin (0.02mg/ml) followed by 4% paraformaldehyde (PFA) in PBS solution. Brains were post-fixed overnight at 4°C then transferred to 30% sucrose solution for 48h. 30μm sections were cut with a cryostat (Leica CM3050S) and a standard free-floating immunohistochemical protocol was used to increase the signal of the tagged ChR2 and counter-stain neurons. The primary antibodies used were anti-GFP (1:1000, chicken; A10262, or rabbit; A11122, Life Technologies), anti-RFP (1:1000, rabbit; 600-401-379, Rockland) and anti-NeuN (1:1000, mouse; MAB-377, Millipore) and the secondary antibodies were Alexa-488 Donkey anti-rabbit and Goat anti-chicken, Alexa-568 Donkey anti-rabbit and Donkey anti-mouse, and Alexa-647 Donkey antimouse (1:1000, Life Technologies). Brain sections were mounted on charged slides using the mounting medium SlowFade Gold (containing DAPI; S36938, Life Technologies), and imaged using a wide-field microscope (Nikon TE2000).

For optogenetic inactivation experiments, VGluT2-Cre and VGluT2::EYFP mice were injected with AAV-DIO-iChloc-dsRed, (see Viruses) into the dmSC (250nl per side, ML: +/− 0.35, AP: 0.1 to −0.45, DV: −1.4 to −1.55) or dPAG (200nl per side, ML: +/− 0.4, AP: −0.4 to −1, DV: −2.2), with 2 injections per hemisphere along the AP axis spaced 300μm apart. Dual optic fibres (400μm diameter, 1.2mm apart, DFC_400/430-0.48_3.5mm_GS1.2_C60; Doric Lenses Inc.) were implanted at the injection site. Behavioural testing was done 10-41days after virus injection. Animals were presented with visual or auditory stimuli that elicited escape, and laser-on trials were interleaved with laser-off trials (473nm, 5-8s square pulse, 15mW/mm^2^). For histological confirmation of the fibre placement and injection site, animals were decapitated under anaesthesia, brains were quickly removed and post-fixed in 4% PFA overnight at 4°C. Slices of 100μm thickness were cut on a HM650V vibratome (Microm) in 0.1M PBS, stained with DAPI before mounting, and imaged on a wide-field microscope (Axio Imager 2, Zeiss).

### Chemogenetic inactivation experiments

VGluT2-Cre and VGluT2::EYFP mice were injected with AAV-DIO-hM4D-nrxn-mCherry (see Viruses) into the dmSC (200-250nl per side, ML: +/− 0.35, AP: −0.1 to −0.45, DV: −1.4 to −1.55), with 2-3 injections per hemisphere along the AP axis. Dual guide cannulae were implanted at ML: +/− 0.6, AP: −0.55, DV: −1.6 to target the SC-dPAG projection, and ML: +/−1.7, AP: +1.7, DV: −2.05 (angle: 7° lateral from zenith) to target the SC-LP thalamus projection. In experiments with optogenetic stimulation, AAV-DIO-ChR2-EYFP was injected into the dmSC first (coordinates and volumes as above) and a 200μm optic fibre cannula was implanted at ML: +/−0.1, AP: −0.3, DV: 1.35 (angle: 35° posterior from zenith). After 20-55days, escape responses to optogenetic or visual stimuli were assessed in a baseline session to estimate the stimulus intensities that evoke escape with P_escape_=1. 30min following microinfusion or i.p. injection, escape responses were reassessed using the same stimuli, and, for optogenetic activation, 200% of baseline intensity or frequency were tested in addition to the baseline strength. For cerebral microinfusions, CNO was diluted in buffered saline containing (in mM): 150 NaCl, 10 D-glucose, 10 HEPES, 2.5 KCl, 1 MgCl2, and to a final concentration of 1 or 10μM. Experiments with visual-evoked escape were done with 1μM, and optogenetically-evoked escape with 1 and 10μM. There was no significant difference between 1 and 10μM at the electrophysiological and behavioural level, and the data have thus been pooled (comparisons between 1μM and 10μM CNO: ChR2-induced firing of SC VGluT2^+^ neurons, P>0.999 Wilcoxon test; SC-dPAG VGluT2^+^ EPSC amplitude, P=0.0973 Mann Whitney test; P_escape_ after CNO microinfusion, P=0.6095, Mann Whitney test). Cerebral microinfusions of CNO or vehicle were performed as described above using 500μm protruding internal cannulae (see Pharmacological Inactivation), with a volume of 0.6-1.0μl per hemisphere. For i.p. injections, 1mg CNO was dissolved in 1ml 0.9% saline just before the experiment and injected at a final concentration of 10mg/kg. Repeated administration of CNO was separated by 2-3 days, preceded by a new baseline session for each treatment. Histological confirmation of cannula placements and viral infection was performed as stated above.

### Electrophysiological recordings in acute midbrain slices

***Data acquisition***: Coronal slices were prepared from VGluT2::EYFP mice aged 6-12 weeks. Brains were quickly removed and transferred to ice-cold slicing solution containing (in mM): 87 NaCl, 26 NaHCO_3_, 50 sucrose, 10 glucose, 2.5 KCl, 1.25 NaH_2_PO_4_, 3 MgCl_2_, 0.5 CaCl_2_. Acute coronal slices of 250μm thickness were prepared at the level of the SC and PAG (−4.8 to −4.1mm from bregma) using a vibratome (VT1200, Leica or 7000smz-2, Campden). Slices were then stored under submerged conditions, at near-physiological temperature (35°C) for 30min before being cooled down to room temperature (19-23°C). For recordings, slices were transferred to a submerged chamber and perfused with ACSF containing (in mM): 119 NaCl, 26 NaHCO_3_, 10 glucose, 2.5 KCl, 2 CaCl_2_, 1 MgCl_2_, 1 NaH_2_PO_4_ (heated to 34°C at a rate of 2-3 ml min^-1^). All ACSF was equilibrated with carbogen (95% O_2_, 5% CO_2_, final pH 7.3). Whole-cell patch-clamp recordings were performed with an EPC 800 amplifier (HEKA). Data was digitised at 20kHz (PCI 6035E, National Instruments), filtered at 5kHz and recorded in LabVIEW using custom software. Pipettes were pulled from borosilicate glass capillaries (Harvard Apparatus, 1.5mm OD, 0.85mm ID) with a micropipette puller (P-1000, Sutter, USA or P-10, Narishige, Japan) to a final resistance of 4-6MΩ. Pipettes were backfilled with internal solution containing (in mM): 130 KGluconate or KMeSO_3_, 10 KCl, 10 HEPES, 5 phosphocreatine, 2 Mg-ATP, 2 Na-ATP, 1 EGTA, 0.5 Na_2_-GTP, 285-290mOsm, pH was adjusted to 7.3 with KOH. VGluT2^+^ dPAG and dmSC cells were visualised on an upright Slicescope (Scientifica) using a 60x objective (Olympus) and identified based on location and EYFP expression. The resting membrane potential (RMP) was determined immediately after establishing the whole-cell configuration and experiments were only continued if cells had a RMP more hyperpolarised than −45mV. Input resistance (R_in_) and series resistance (R_s_) were monitored continuously throughout the experiment, and Rs was compensated in current-clamp recordings. Only cells with a stable R_s_ <30MΩ were analysed. For ChR2-assisted circuit mapping, recordings were made 10-51 days (mean= 22.3±2.3 days) after injection of AAV2-DIO-ChR2-mCherry into the mSC or dPAG of VGluT2::EYFP mice. ChR2 was stimulated with wide-field 490nm LED illumination (pe-100, CoolLED, 1ms or 10ms pulse length, maximum light intensity=2.7mW). To characterise the cellular effects of iChloc activation, dPAG or dmSC VGluT2^+^ cells expressing AAV5-DIO-iChloc-dsRed were recorded from at 46 days after infection, and iChloc was stimulated with 1s long, 490nm light pulses. Recordings in animals expressing hM4D-nrxn in the dmSC were made 22-53 days after injection (mean= 29.4±3.1 days), and ChR2 was activated at 10, 20 and 100% light intensity (0.27, 0.54 and 2.7mW).

***Pharmacology***: No drugs were added to the recording ACSF, except for the following experiments: miniature EPSCs (mEPSCs) were recorded in 1μM Tetrodotoxin (TTX, Sigma Aldrich), and ESPC recordings shown in Figure S10 were recorded in 1μM TTX and 100μM 4-Aminopyridine (4-AP, Sigma Aldrich); to test the effect of hM4D-neurexin activation on firing rates and synaptic transmission, 1-10μM CNO (freebase, Hellobio) was added to the ACSF during recordings.

***Data analysis***: Analysis was performed using custom-written procedures in Python, except for the analysis of sEPSCs and mEPSCs which was done in IGOR Pro 6 (WaveMetrics) using TaroTools (by Taro Ishikawa). The R_in_ was calculated from the steady-state voltage measured in response to a hyperpolarising test pulse of 500ms duration at a holding potential of −60mV. The membrane time constant was calculated by fitting the decay of the test pulse with a single exponential (y=*y*0+A*e*^(-(*x-x*0)/*τ*)^). The membrane potential values stated in the text are not corrected for liquid junction potentials. The sEPSC frequency before and after ChR2 stimulation was calculated from 6-8 repetitions per cell. Failures of light-evoked synaptic transmission were defined as a peak amplitude of less than the mean current baseline +2SD in a time window defined by the onset of the mean evoked synaptic current ±5ms. Quantal content calculated by the direct method^41,42^ was obtained by dividing the peak amplitude of the evoked current by the peak amplitude of the sEPSCs in the same cell (which is not significantly different from the mEPSC amplitude, see Figure S11). The paired-pulse ratio was calculated as the ratio of peak amplitudes between the second and first EPSCs in a train. Effects of drug application were calculated after a perfusion time of at least 10min. Statistical analysis was performed on cells pooled across animals.

### Single unit recordings

***Data acquisition***: Neuropixels silicon probes (phase3, option1, 384 channels^46^) were used to record extracellular spikes from dmSC neurons in three male adult C57BL/6J wild-type mice. A craniotomy was made over the SC and sealed with Kwik-Cast, followed by attachment of a metal custom-made head-plate and ground pin to the skull, using dental cement. At least 36 hours after surgery, mice were placed on a plastic wheel and head-fixed at an angle of 30° from the anterior-posterior axis, parallel to an LCD monitor (Dell, 60Hz refresh rate) centered 30 cm above the head. Prior to recording, the probe was coated with DiI (1mM in ethanol, Invitrogen) for track identification and a wire was connected to the ground pin for external reference and ground. For recording, the probe was slowly inserted into the SC (AP: −0.5 to −0.7, ML: 0.4 to 0.8) to a depth of 2.8-3.0 mm and left in place for at least 20 minutes before the beginning of the recording session. Data was acquired using spikeGLX (https://github.com/billkarsh/SpikeGLX, Janelia Research Campus), high-pass filtered (300Hz), amplified (500x), and sampled at 30kHz. Sensory stimuli were delivered and synchronized using custom-made LabVIEW software and a PCIe-6353 board (National Instruments). Visual and auditory stimuli (98% contrast; 50% contrast; sound) were presented interleaved with a 1min interval and a total of 30 presentations each.

***Data Analysis***: Analysis was performed in MATLAB 2017a. Raw voltage traces were band-pass filtered (300-5000Hz), spikes were detected and sorted automatically using JRCLUST^70^, followed by manual curation. Only units with a clear absolute refractory period in the auto-correlogram were classified as single units. Firing rate histograms were calculated as the average firing rate in bins of 1ms for 30 consecutive trials, and subsequently smoothed. Units were considered to respond to the threat stimulus if their firing rate increased by at least 1Hz in a 500ms time-window from stimulus onset when compared to the baseline (500ms before stimulus onset). Peak firing rates for each stimulus were calculated as the mean of a 30ms time-window centered on the time of the average peak firing rate of all responding units. Responses to 50% contrast visual stimuli were calculated on all units that responded to 98% contrast. For units showing persistent activity after stimulus offset, the time constant to decay to baseline was obtained by fitting a single exponential to the average firing rate histogram. Statistical analysis was performed on single units pooled from all animals.

**Retrograde Tracing**: For monosynaptic rabies tracing from the dPAG, TVA and RG were injected unilaterally into the dPAG with an angled approach from the contralateral hemisphere to avoid infection of the SC in the target hemisphere (20°, AP: −0.45 to −0.5, ML: −0.6, DV: −2.2,). EnvA-dG-RV-mCherry was injected into the dPAG vertically (AP: −0.4, ML: +0.5, DV: −2.1) 10-14days later. Animals were perfused seven days post-rabies virus injection. Brains were cut at 100μm thickness on a microtome (HM650V, Microm). All sections containing the PAG and SC were mounted in SlowFade Gold, and imaged using a confocal microscope (SP8, Leica). Tile scans of the entire section were acquired with a 25x water objective (Olympus) at five depths (10μm apart) and maximum projections of these stacks were used for subsequent analysis. Cell counting was done manually (Cell counter plug-in, Fiji) in reference to the Paxinos and Franklin atlas^71^. To quantify the position of presynaptic SC cells along the mediolateral axis, the coordinates of the counted cells were normalised to the medial and lateral extents of the SC for each brain slice, and a kernel density estimation was performed (Scikit-learn, Python). For retrograde tracing from the dmSC, rAAV2-retro-mCherry was injected unilaterally. AAV2-CamkII-GFP was co-injected to label the injection site in 2 out of 3 brains. Animals were sacrificed 14-18 days afterwards and their brains processed as described above. Every third section along the rostrocaudal axis of the SC was imaged with on an Axio Imager 2 (Zeiss) and presynaptic cells in the dPAG and auditory cortex were counted manually.

**Histological quantifications**: To estimate the fraction of VGluT2^+^ cells in a target area that were infected with viral vectors, we compared the density of infected cells in VGluT2^Cre^ animals at the implant site, to control densities quantified using the VGluT2::EYFP reporter line. Optogenetic vectors infected 86±6% for dPAG and 95±9% for mSC; GCamp6 infected 90±8% for dPAG and 86±1% for mSC; hM4D infected 93±15% for mSC.

**General Data analysis**: Data analysis was performed using custom-written routines in Python 2.7. and custom code will be made available per request. Data are reported as mean±SEM unless otherwise indicated. Statistical comparisons using the significance tests stated in the main text were made in SciPy Stats and GraphPad Prism, and statistical significance was considered when P<0.05. Data were tested for normality with the Shapiro-Wilk test, and a parametric test used if the data were normally distributed, and a non-parametric otherwise, as detailed in the text next to each comparison.

## Acknowledgments

This work was funded by a Wellcome Trust/Royal Society Henry Dale Fellowship (098400/Z/12/Z), a Medical Research Council (MRC) grant MC-UP-1201/1, a Wellcome Trust and Gatsby Charitable Foundation SWC Fellowship (to T.B.), MRC PhD Studentship (to D.E. and R.V.), a Boehringer Ingelheim Fonds PhD fellowship (to R.V.), DFG fellowship (to A.V.S and S.R.), and a Marie Skłodowska-Curie Individual Fellowship (706136) and EMBO Long Term Fellowship (to Y.L). We thank P. Latham and members of the Branco lab for discussions, S. Sternson, P. Dayan, T. Margrie and T. Mrsic-Flogel for comments on the manuscript, S. Sternson, S.Wiegert, T.Oertner, T.Margrie for generous gifts of viral vectors, P. Iordanidou, T. Okbinoglu and L. Jin for technical assistance, D.Campagner, T.Harris and N.Steinmetz for help with silicon probe recordings, and K. Betsios for programming the data acquisition software.

